# Stimjim: open source hardware for precise electrical stimulation

**DOI:** 10.1101/757716

**Authors:** Nathan Cermak, Matthew A. Wilson, Jackie Schiller, Jonathan P. Newman

## Abstract

Electrical stimulation is a simple and powerful tool to perturb and evoke neuronal activity in order to understand the function of neurons and neural circuits. Despite this, devices that can provide precise current or voltage stimulation are expensive and closed-source. Here, we introduce Stimjim, a capable and inexpensive ($200 USD) open-source instrument for electrical stimulation that combines both function generation and electrical isolation. Stimjim provides microsecond temporal resolution with microampere or millivolt scale precision on two electrically isolated output channels. We demonstrate Stimjim’s utility both *in vitro* by precisely stimulating brain slices, and *in vivo* by training mice to perform intracranial self-stimulation (ICSS) for brain stimulation reward. During ICSS, Stimjim enables the experimenter to smoothly tune the strength of reward-seeking behavior by varying either the output frequency or amplitude. We envision Stimjim will enable new kinds of experiments due to its open-source and scalable nature.

## Introduction

Electrical stimulation of neural tissue is an invaluable and ubiquitous research tool. Over the past 150 years, it has helped researchers understand the function of various brain regions by directly inducing neurons in those regions to fire ^1;2;3^. More recently, it has also found important clinical applications in neurological disorders including Parkinson’s disease ^4^ and depression ^5^. However, to date, the hardware for performing precise current- and voltage-based electrical stimulation generally remains expensive and closed source.

In contrast, there has been a recent push within the scientific community to produce open labware – open source hardware and software replacements for a variety of common laboratory tasks ^6;7;8^. Examples in the life sciences include software and hardware for:

- recording or stimulating neurons (e.g. Open Ephys ^9;10^, Miniscopes ^11;12;13;14^, and others ^15;16;17^)
- amplifying DNA (e.g., OpenPCR ^18^)
- fluid control^19^ and turbidostats^20;21;22^
- microscopy ^23;24;25;26^ and microscope components^27^
- plate readers and spectrophotometers ^28;29^
- electroporation ^30^
- ecological monitoring (e.g., Audiomoth ^31^)

We now add Stimjim to this growing body of open hardware. Stimjim replaces commercial neural stimulators at a fraction of the cost, with improved programmability. Furthermore, due to its entirely open design and software, Stimjim can be modified by users to fit their specific needs.

## Results

### Design

We developed Stimjim to be a precise, electrically isolated stimulus generator. Stimjim is based on the Teensy 3.5 microcontroller board (www.pjrc.com/teensy), which utilizes a 32-bit Arm Cortex-M4F processor running at 120 MHz. Each stimulating channel includes a current source based on an improved Howland current pump ^32^, and a voltage source (an op-amp), driven by a 16-bit digital-to-analog converter (DAC). The final output of each channel is selected by a 4-way switch, such that either channel can be configured as a current output, voltage output, grounded, or disconnected. To ensure the stimulator is properly connected (a common issue with experiments in freely moving animals) and to verify required stimulus current or voltage amplitudes, each channel also has an analog-to-digital converter (ADC) able to read either the output voltage or the output current (via a low-value sense resistor in series with the current output). Our circuit board design was made using Kicad ^33^ (www.kicad-pcb.org), an open-source printed circuit board (PCB) design program. Schematic, layout, bill of materials, and build instructions are included as supplemental materials and are also available in the Stimjim git repository (bitbucket.org/natecermak/stimjim).

Stimjim’s design compares favorably against alternatives (Table 1). It is an order of magnitude less expensive than most commercial alternatives. Its only draw-back is that its compliance voltage is lower, which limits the load resistance that Stimjim can drive. For a given resistance *R*, each Stimjim channel cannot output a current larger than 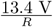. Thus, Stimjim is not suitable for high-impedance electrodes such as pulled glass electrodes. Note however, that Stimjim’s compliance voltage can be doubled to roughly ± 27 V by connecting the two channels in series. As a low-cost open-source device, Stimjim is perhaps most similar to PulsePal 2, an open-source programmable pulse generator ^34^. However, it expands on PulsePal’s capabilities by including electrical isolation, current output mode, and on-board monitoring of output currents/voltages. Further electrical characteristics of Stimjim are given in Table 2.

**Table 1:** Comparison of Stimjim vs other related hardware. Cost was obtained as either the complete cost of the bill of materials (Stimjim and PulsePal) or quoted prices from distributors (STG400x, PHM-15x, ISO-01B). ^*a*^ISO-01B has a compliance warning sound if the load resistance is too high, but does not report actual current measurements. ^*b*^Compliance voltage can be doubled by connecting both output channels in series.

|  | Stimjim | PulsePal 2 | STG400x<br>(Multichannel Systems) | PHM-15x<br>(Med Associates) | ISO-01B<br>(NPI) |
| --- | --- | --- | --- | --- | --- |
| Output channels | 2 | 4 | 2-8 | 2 | 1 |
| Stimulus generator included | Y | Y | Y | Y | N |
| Current output mode | Y | N | Y | Y | Y |
| Voltage output mode | Y | Y | Y | N | Y |
| Outputs electrically isolated | Y | N | Y | Y | Y |
| Onboard measurement | Y | N | N | Optional | Y<sup>a</sup> |
| Compliance voltage | $\pm 15$ V (voltage)<br>$\pm 13.7$ V<sup>b</sup> (current) | $\pm 10$ V | $\pm 120$ V | $\pm 45$ V | $\pm 100$ V |
| Fastest pulse | 20 μs | 100 μs | 20 μs | 60 μs | 10 μs |
| Cost (USD) | \$202 (parts) | \$264 (parts)<br>\$745 (assembled,<br>from Sanworks) | \$4131 (2 channels)<br>\$7462 (4 channels)<br>\$10327 (8 channels) | \$6211 (+ \$4438<br>for software) | \$1708 |
| Open source | Y | Y | N | N | N |

**Table 2:** Stimjim electrical characteristics. Parameters were measured on Instek GDS-1054B digital oscilloscope, full bandwidth (50 MHz). ^*a*^Calculated for 100 ms segments. Note that this is the noise only when the pulse is delivered; at all other times the output is directly connected to ground. Voltage noise in current mode was measured with a 9.86 kΩ resistive load. ^*b*^Calculated as output range divided by resolution of output DAC (16 bit). ^*c*^value from from datasheet for Vishay DG509B (output switch).

|  | Value | Units |
| --- | --- | --- |
| Slew rate (voltage mode) | 7.2 | V$\cdot\mu$s<sup>-1</sup> |
| Slew rate (current mode) | 3 | V$\cdot\mu$s<sup>-1</sup> |
| Output voltage noise (voltage mode)<sup>a</sup> | 0.8 | mV rms |
|  | 7 | mV p-p |
| Output voltage noise (current mode)<sup>a</sup> | 0.6 | mV rms |
|  | 5 | mV p-p |
| Min. voltage increment<sup>b</sup> | 0.45 | mV |
| Min. current increment<sup>b</sup> | 0.1 | μA |
| Output impedance (voltage mode)<sup>c</sup> | 180 | $\Omega$ |
| Trigger latency | 10 | μs |

Stimjim’s software is written in C++ using the the Arduino development environment. We provide an Arduino-compatible Stimjim library permitting low-level device control (writing registers in the DACs or ADCs, or setting the stimulation control mode). Library functions enable users to create new programs to run on Stimjim for example, generation of custom waveform outputs stored on the onboard SD card. We also provide a default program using this library that can generate user-defined pulse train sequences. Users set the parameters for pulse trains and read the measured pulse amplitudes via a 12 Mbit/s serial connection over USB. Pulse train parameters include output mode (current or voltage), frequency, duration, and the amplitude of each phase of the pulse itself. Stimjim can store definitions for 100 pulse trains concurrently, and users can select and initiate particular pulse trains on the fly.

### Benchmarking

To benchmark Stimjim and our pulse train program, we generated a series of one-second biphasic pulse trains in which we varied the pulse frequency (from 2 Hz to 4000 Hz), pulse duration (from 20 *µ*s to 4000 *µ*s), and amplitude. We simultaneously recorded from both of Stimjim’s output channels using a National Instruments PCI-6110 card (2 MHz sampling rate per channel, 4.9 mV resolution). One Stimjim channel was set to voltage mode and the other channel to current mode with a 9.86 kΩ resistor connected to the output.

Stimjim proved capable of providing microsecond temporal resolution and millivolt- and microampere-amplitude resolution. Across the tested range of stimulation frequencies, Stimjim generated accurate and highly consistent inter-pulse intervals (IPIs; Fig. 2A-C) and pulse widths (PW) (Fig. 2D-F). While worst case errors of 2 *µ*s (IPI) and 10 *µ*s (PW) were detected, typical performance exceeded the temporal resolution of our test equipment. For example, IPI and PW standard deviations were typically less than 0.5 *µ*s, which was the temporal resolution of our test equipment. For both IPI and PW, the absolute error magnitudes increased as the duration itself increased (Fig. 2C,F). However, the worst case absolute errors (2 *µ*s and 10 *µ*s) correspond to fractional errors of 0.0004% and 0.25% for IPI and PW, respectively. Finally, we assessed pulse amplitudes across a range of settings to ensure negligible DC offsets and proper gains. From −10V to +10V (the range of our test equipment), Stimjim produced accurate voltage and current amplitudes, with maximal errors of less than 40 mV and 2.5 *µ*A (Fig. 2G-I). Pulse rise and fall times were rapid (Fig. 2J and Table 2) and exhibited low noise (Table 2). However, we did observe small-amplitude (0.2 V) high-frequency spikes during voltage pulses, which resulting from reading the output voltage via the onboard ADC. If needed, users can remove the ADC read operation and eliminate these spikes. Current pulses did not exhibit such spikes because the ADC instead reads a buffered signal from the current-sense amplifier, not the actual output signal.

**Figure 1:**
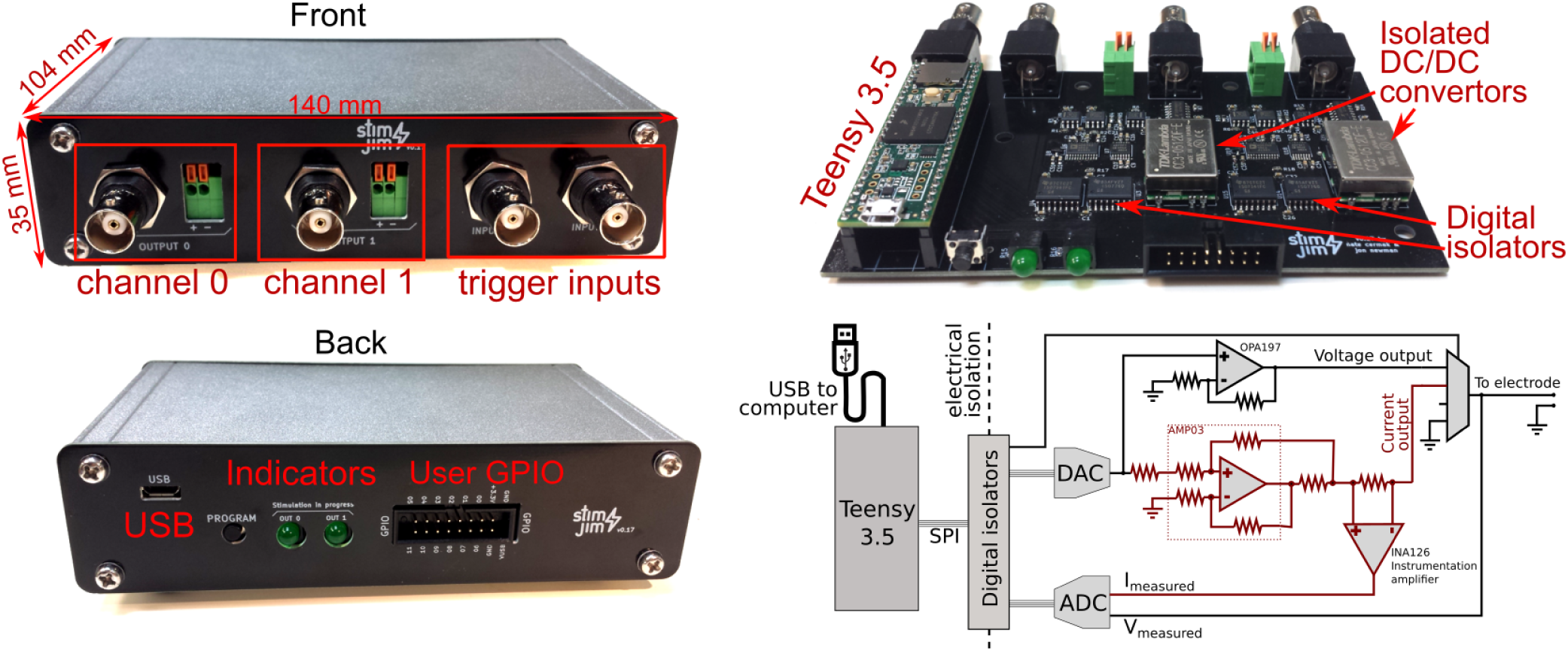
Stimjim is a compact, configurable, and precise stimulator. Stimjim has a compact footprint, measuring 104 × 140 × 35 mm. The front panel (top left) includes BNC and push-terminal connectors for the outputs, and BNC connectors for inputs. On the back (lower left) are the USB connector (which also provides power), LED indicators for active stimulation, and a set of breakout pins for general purpose user input/output (GPIO). While we do do not demonstrate using these GPIO pins in this paper, we provide them for advanced users who may want them. The internal circuit board (top right) consists of a Teensy 3.5 and two electrically isolated output channels. Each output channel has its own isolated DC-DC power convertor and high-speed digitial isolators for communicating with the Teensy. The lower-left panel shows the basic circuit for each channel. A digital-to-analog convertor (DAC) provides the analog signal to both the current and voltage output circuits. The voltage output circuit consists of a non-inverting amplifier (OPA197 op-amp) with a gain of 1.5. The current output circuit is marked with red wires, and uses a difference amplifier (AMP03, which includes four internal 25 kΩ laser-trimmed resistors), with two external 3 kΩ 0.1% resistors. The current output circuit includes a small series resistor that enables measuring the output current with an onboard analog-to-digital convertor (ADC). A 4-way switch enables selecting the voltage output, current output, or grounding or disconnecting the output. The ADC can also measure the voltage at the output terminal.

**Figure 2:**
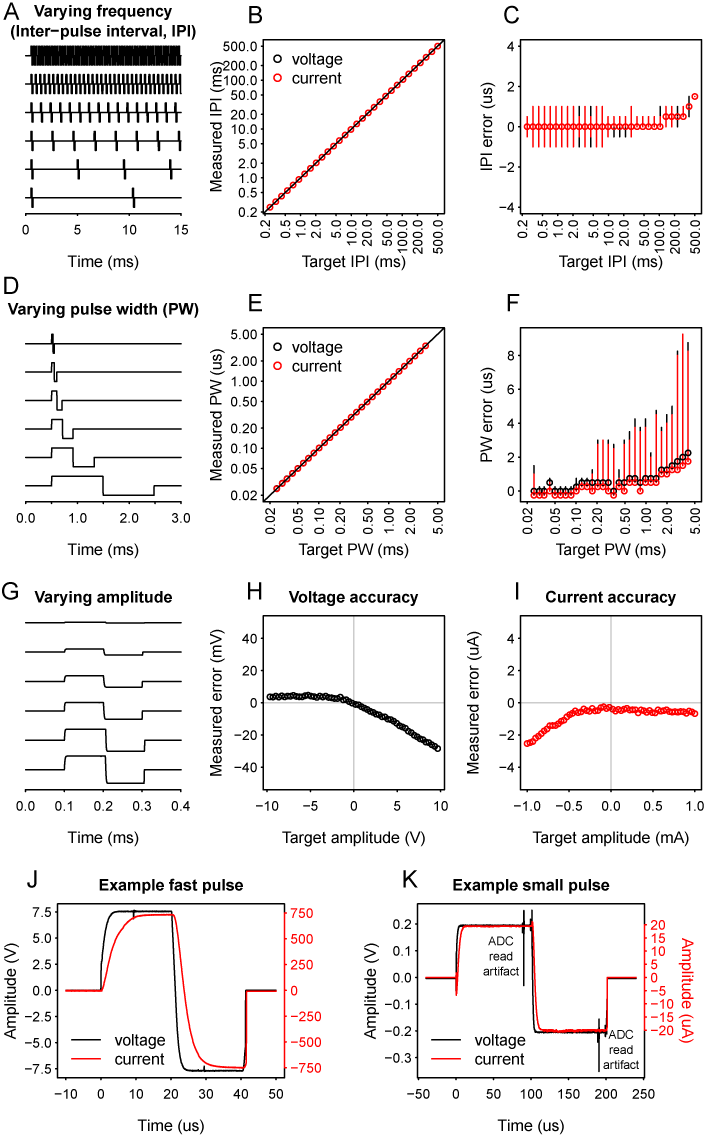
Electrical benchmarks show Stimjim provides microsecond temporal precision and millivolt- and microampere-scale amplitude precision. (A) Example pulse trains with varying inter-pulse interval (IPI). (B) Median IPI in a one-second pulse train measured on a high-speed data acquisition device, plotted against target IPI. Red dots show current output, black dots show voltage output. Solid line shows equality. (C) Plot of IPI errors measured in a 1-second pulse train. Points show median error, error bars indicate worst-case errors. (D) Example pulse trains with varying pulse width (PW). (E) Median measured PW vs target PW. Red dots show current output, black dots show voltage output. Solid line shows equality. (F) Plots of pulse width errors over 100 pulses. Conventions are the same as for (C). (G) Example pulses with varying amplitude. (H) Error in amplitude of voltage pulses (100 *µ*s, 1 kHz) vs target amplitude. (I) same as (H) but for current pulses. (J) Example of a fast biphasic (20 *µ*s/phase) pulse. Red line is a 750 *µ*A current pulse (with a 9.86 kΩ resistive load), black line is a 7.5 V pulse. (K) Example of a small-amplitude pulse. Red line is a 20 *µ*A current pulse (with a 9.86 kΩ resistive load), black line is a 0.2 V pulse. Pulses shown (J) and (K) were measured on oscilloscope for higher bandwidth and reduced input capacitance.

### Brain slice stimulation

We evaluated Stimjim for use in brain slice experiments. While Stimjim could not provide sufficient current for synaptic stimulation through pulled glass theta electrodes (resistance greater than 1 MΩ, data not shown), we were able to successfully stimulate pyramidal neurons in rat piriform cortex slices using monopolar platinum-iridium electrodes (100 kΩ). The exposed conical electrode tip was approximately 20 *µ*m long with a maximal diameter of roughly 5 *µ*m. We first placed a single stimulating electrode approximately 100 *µ*m adjacent to the soma amd applied 0.4 ms current stimulation pulses of gradually increasing amplitudes. With increasing amplitude, we observed increasingly rapid and reliable action potential generation, and eventually emergence of a second action potential (Fig. 3A).

**Figure 3:**
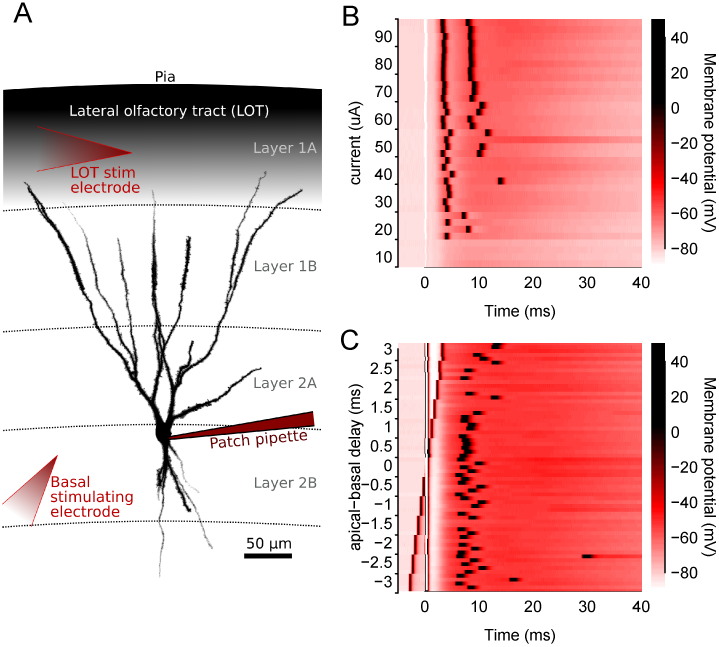
Stimjim can be used for brain slice stimulation. (A) Pyramidal neuron in piriform cortex filled with CF633, showing locations of stimulating electrodes (red arrows) for cell recorded in (C). Setup for (B) was similar but with a different cell and without the LOT stimulating electrode. (B) Increasing stimulus intensity through an electrode positioned adjacent to basal dendrites of a pyramidal cell first variably evokes an action potential, and then eventually a second action potential. (C) Stimjim’s two channels can be used to provide precise time-delays between different stimulus electrodes. This neuron was stimulated via an electrode positioned in the lateral olfactory tract (LOT) and another electrode adjacent to soma and basal dendrites. Trials are aligned to the stimulus artifact from the basal electrode.

Next, we verified Stimjim’s ability to provide coordinated pulses on two separate electrodes. We placed one electrode in the lateral olfactory tract (LOT), a thick layer of axons that courses through the apical dendrites of piriform pyramidal neurons. We then placed a second electrode approximately 100 *µ*m from the soma, near the basal dendrites. We generated variable delays (up to ± 3 ms) between LOT and basal stimulation (Fig. 3B). When LOT inputs were stimulated 0.5 ms after basal stimulation, but not before, we observed the most reliable generation of action potential. Outside of this window, action potential timing was variable and action potentials occasionally were not evoked. These experiments demonstrate Stimjim’s potential for precise extracellular electrical stimulation in brain slices.

### *In vivo* stimulation

To demonstrate Stimjim’s utility *in vivo*, we used it to train mice in a classical paradigm known as intra-cranial self stimulation (ICSS) ^35^. In this assay, animals are implanted with electrodes (or more recently optical fibers ^36;37;38^) enabling activation of a pleasure/reward-related brain region ^39^. Animals are then placed in a training paradigm in which they learn that a simple motor action (typically spinning a wheel or pressing a lever) causes direct activation of this brain region. Animals quickly learn the required action and are willing to repeat it for extended periods of time.

We trained two mice in a head-fixed variant of ICSS, in which animals could lick a sensor in order to obtain brain stimulation reward (BSR). We used a capacitive sensor attached to a small metal pole to detect licking, and every lick triggered a stimulus pulse train (0.5 seconds, initially 150 Hz and the minimal current at which animals would respond). To initially encourage licking, we placed a small amount of peanut butter on the metal sensor. After initial licking was reinforced by BSR, animals would continue licking long after the peanut butter was gone, including during the next session in which no peanut butter was offered. After animals had learned the licking behavior (usually within their first hour session), we varied the BSR frequency and amplitude and assessed how it affected licking behavior. Both animals showed clear frequency- and amplitude-dependent responses, in which animals ceased licking when the rewarding stimulation was insufficiently intense (Fig. 4).

**Figure 4:**
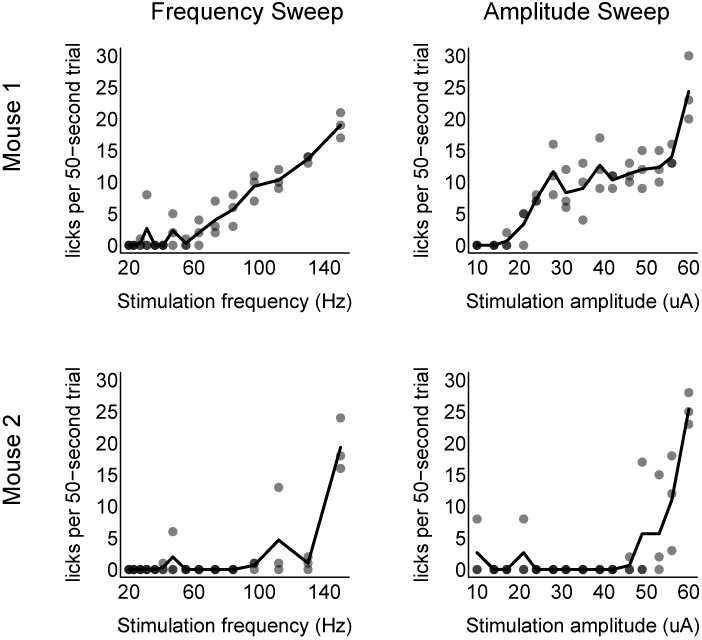
Stimjim enables measuring frequency- and amplitude-dependent responses in an intracranial self-stimulation paradigm. Mice decreased their licking rates when the frequency and amplitude of the rewarding stimulation decreased. For each animal, three frequency sweeps (highest to lowest frequency, one minute per frequency) were performed in a 45-minute session. Amplitude sweeps were performed in the same way. Each dot indicates a single one-minute trial, and the solid black line shows the mean of all three trials at that frequency or amplitude.

We observed clear differences between the two animals. Mouse 1 shows a rather linear response to either increasing frequency or increasing amplitude, whereas mouse 2 had a more “digital” response akin to passing an activation threshold. However, maximal licking rates were comparable between the two animals. Such differences are likely due to electrode placement^35^, although they may also reflect intrinsically different personalities between the two animals. Stimjim provides a precise and cost-effective means to scan the space of stimulation patterns, which could be useful to ensure all animals are given stimuli yielding the same response level.

As a secondary test of Stimjim’s ability to provide effective BSR, we placed head-fixed mice on a linear treadmill and recorded their running behavior for 20 minutes. We then offered BSR for every increment the mice ran on the treadmill, initially every 20 cm and linearly increasing up to 60 cm over the course of 20 minutes. As shown in Fig. 5, mice always ran faster when BSR was offered than when it was not (n=9 sessions across 4 mice, p=0.004, paired Wilcoxon rank sum test). This shows that Stimjim provides a cost-effective means of motivating mice to run, such as for experiments studying place cells or motor-related neural signaling.

**Figure 5:**
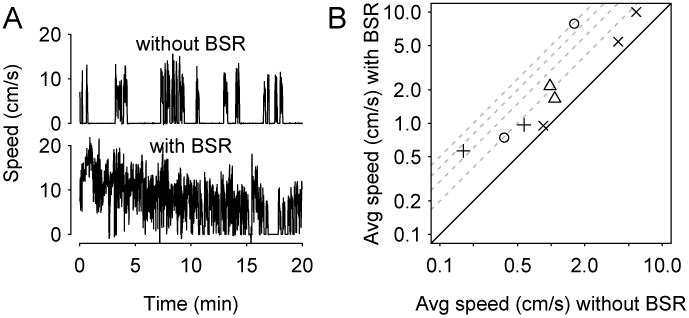
Stimjim can provide brain stimulation reward to encourage head-fixed running behavior. Head-fixed mice on a 1-D treadmill significantly increased their running when given BSR for every 20-60 cm they moved. (A) Example mouse showed occasional running bouts without any reward (top). However, when BSR was offered for every 20 cm the mouse ran (and gradually increased to every 60 cm by the end of the experiment), the mouse ran consistently for nearly the entire 20 minutes. (B) Speed was always higher when reward was offered, across four mice (symbols) and 2-3 sessions per mouse. Solid line shows equality, dashed grey lines show 2-, 3-, 4- and 5-fold increases.

## Conclusions

We have introduced Stimjim, an inexpensive yet precise open-source stimulator for neuroscience. At a cost of roughly $200 USD for parts, Stimjim is order of magnitude less expensive than commercial, proprietary alternatives. It offers microsecond temporal control of current and voltage with millivolt/microampere precision.

Stimjim’s low cost opens up many potential applications, such as learning paradigms that involve direct electrical stimulation. Stimjim’s open source nature makes it straightforward for researchers to customize the stimulation parameters and use Stimjim in closed loop experiments. Furthermore, researchers who were previously limited to training only one animal at a time due to possessing only a single stimulator could now train or perform experiments with ten or more animals simultaneously for comparable cost.

## Supporting information

Fabrication (Gerber) files for main board

Fabrication (Gerber) files for enclosure panels

Bill of materials

Stimjim circuit schematic

## Acknowledgments

We thank Amit Kumar for help with the brain slice experiments, and Jakob Voigts for support in the MFB implant surgery. We also thank the Schiller lab for discussions and help with preliminary testing. NC acknowledges support in part from a Zuckerman STEM Leadership Fellowship at the Technion.

## Methods

### Stimjim fabrication and benchmarking

PCBs for Stimjim were ordered from JLCPCB and components were ordered from Digikey. Components were manually soldered to the PCB using solder paste and a soldering iron. After soldering, the pulse control program was downloaded to the Teensy using the Arduino IDE and Teensyduino. From that point on, Stimjim’s settings were controlled via serial communication over USB. For benchmarking, we used a custom NI LabView program to set Stimjim’s pulse parameters (frequency, amplitude, duration, etc.), initiate a one-second pulse train, and record both Stimjim channels using a National Instruments PCI-6110 card via a breakout box. This program is also available in the git repository.

### Electrode implantation and ICSS

Monopolar electrodes (Plastics1, #MS303/2-AIU/SPC, coated stainless steel, 200 *µ*m diameter) were implanted above the medial forebrain bundle according to the protocol in reference ^39^. The ground was implanted in the contralateral cortex. Additionally, a 3D-printed headpost was affixed to the animal’s skull by dental cement to enable head fixation. Typical resistance (100 *µ*s pulse) between connector pins after implantation was 20-30 kΩ. All animal procedures were in accordance with guidelines established by the NIH on the care and use of animals in research and were confirmed by the Technion Institutional Animal Care and Use Committee (IL-012-01-18, valid until 10/4/2022).

### Slice stimulation experiments

Coronal brain slices were prepared from the anterior piriform cortex from 28-40 day old Wistar rats. 300 *µ*m thick slices were cut in ice-cold artificial cerebro-spinal fluid (ACSF) bubbled with 95% oxygen and 5% CO_2_, then incubated for 30 min at 37 C and kept at room temperature afterwards. Whole cell patch clamp recordings were performed with an Axon amplifier (Multiclamp). Glass electrodes (6-8 MΩ) were made from thick-walled (0.25 mm) borosilicate glass capillaries on a Flaming/Brown micropipette puller (P-97; Sutter Instrument). Intracellular pipette solution contained 135 mM potassium gluconate, 4 mM KCl, 4 mM Mg-ATP, 10 mM Na_2_-phosphocreatine, 0.3 mM Na-GTP, 10 mM HEPES, 0.2 mM OGB-6F, 0.2 mM CF-633, and biocytin (0.2%) at pH 7.2. The ACSF solution contained 125 mM NaCl, 25 mM NaHCO_3_, 25 mM Glucose, 3 mM KCl, 1.25 mM NaH_2_PO_4_, 2 mM CaCl_2_, 1 mM MgCl_2_ at pH 7.4. After patches were established, platinum-iridium electrodes for stimulation (Alpha Omega, #387-102S01-11, 250 *µ*m diameter, Parylene C and Polyamide coated, 0.1 MΩ) were placed in the lateral olfactory tract and in the basal dendrites roughly 100 *µ*m from the soma.

## Supplemental materials

In case of future modifications, the most up-to-date details regarding Stimjim will be available at https://bitbucket.org/natecermak/stimjim. The following are included as supplemental materials for the version of Stimjim documented here (v0.18).

- Bill of materials: stimjim bom.xlsx
- Fabrication files: stimjimFabricationFiles v0.18.zip and stimjimPanelFabricationFiles v0.18.zip
- Schematic: schematic.pdf

## References

[1] Eduard Hitzig and G. T. Fritsch. Über die elek trische Erregbarkeit des Grosshirns. Arch Anat Physiol, pages 300–332, 1870.

[2] David Ferrier. The Functions of the Brain: With numerous illustrations. Smith, 1876.

[3] Wilder Penfield and Edwin Boldrey. Somatic motor and sensory representation in the cerebral cortex of man as studied by electrical stimulation. Brain, 60(4):389–443, 1937.

[4] Günther Deuschl, Carmen Schade-Brittinger, Paul Krack, Jens Volkmann, Helmut Schfer, Kai Btzel, Christine Daniels, Angela Deutschlnder, Ulrich Dillmann, Wilhelm Eisner, Doreen Gruber, Wolfgang Hamel, Jan Herzog, Rüdiger Hilker, Stephan Klebe,Manja Klo, Jan Koy, Martin Krause, Andreas Kupsch, Delia Lorenz, Stefan Lorenzl, H. Maximilian Mehdorn, Jean Richard Moringlane, Wolfgang Oertel, Marcus O. Pinsker, Heinz Reichmann, Alexander Reu, Gerd-Helge Schneider, Alfons Schnitzler, Ulrich Steude, Volker Sturm, Lars Timmermann, Volker Tronnier, Thomas Trottenberg, Lars Wojtecki, Elisabeth Wolf, Werner Poewe, and Jürgen Voges. A Randomized Trial of Deep-Brain Stimulation for Parkinson’s Disease. New England Journal of Medicine, 355(9):896–908, August 2006.

[5] Helen S. Mayberg, Andres M. Lozano, Valerie Voon, Heather E. McNeely, David Seminowicz, Clement Hamani, Jason M. Schwalb, and Sidney H. Kennedy. Deep Brain Stimulation for Treatment-Resistant Depression. Neuron, 45(5):651–660, March 2005.

[6] Joshua M. Pearce. Open-Source Lab: How to Build Your Own Hardware and Reduce Research Costs. Elsevier, 1st edition, October 2013.

[7] Tom Baden, Andre Maia Chagas, Greg Gage, Timothy Marzullo, Lucia L. Prieto-Godino, and Thomas Euler. Open Labware: 3-D Printing Your Own Lab Equipment. PLoS Biology, 13(3), March 2015.

[8] Joshua H. Siegle, Gregory J. Hale, Jonathan P. Newman, and Jakob Voigts. Neural ensemble communities: open-source approaches to hardware for large-scale electrophysiology. Current Opinion in Neurobiology, 32:53–59, June 2015.

[9] Joshua H. Siegle, Aarn Cuevas Lpez, Yogi A. Patel, Kirill Abramov, Shay Ohayon, and Jakob Voigts. Open Ephys: an open-source, plugin-based platform for multichannel electrophysiology. Journal of Neural Engineering, 14(4):045003, June 2017.

[10] Alessio Paolo Buccino, Mikkel Elle Lepperd, Svenn-Arne Dragly, Philipp Hfliger, Marianne Fyhn, and Torkel Hafting. Open source modules for tracking animal behavior and closed-loop stimulation based on Open Ephys and Bonsai. Journal of Neural Engineering, 15(5):055002, July 2018.

[11] Denise J. Cai, Daniel Aharoni, Tristan Shuman, Justin Shobe, Jeremy Biane, Weilin Song, Brandon Wei, Michael Veshkini, Mimi La-Vu, Jerry Lou, Sergio Flores, Isaac Kim, Yoshitake Sano, Miou Zhou, Karsten Baumgaertel, Ayal Lavi, Masakazu Kamata, Mark Tuszynski, Mark Mayford, Peyman Golshani, and Alcino J. Silva. A shared neural ensemble links distinct contextual memories encoded close in time. Nature, 534(7605):115–118, May 2016.

[12] William A. Liberti Iii, L. Nathan Perkins, Daniel P. Leman, and Timothy J. Gardner. An open source, wireless capable miniature microscope system. Journal of Neural Engineering, 14(4):045001, May 2017.

[13] Andres de Groot, Bastijn J. G. van den Boom, Romano M. van Genderen, Joris Coppens, John van Veldhuijzen, Joop Bos, Hugo Hoedemaker, Mario Negrello, Ingo Willuhn, Chris I. De Zeeuw, and Tycho M. Hoogland. NINscope: a versatile miniscope for multi-region circuit investigations. bioRxiv, page 685909, July 2019.

[14] Daniel Aharoni, Baljit S. Khakh, Alcino J. Silva, and Peyman Golshani. All the light that we can see: a new era in miniaturized microscopy. Nature Methods, 16(1):11–13, January 2019.

[15] Jonathan P. Newman, Jakob Voigts, Maxim Borius, Mattias Karlsson, Mark T. Harnett, and Matthew A. Wilson. Twister3: a simple and fast microwire twister. bioRxiv, page 727644, August 2019.

[16] Jonathan P Newman, Ming-fai Fong, Daniel C Millard, Clarissa J Whitmire, Garrett B Stanley, and Steve M Potter. Optogenetic feedback control of neural activity. eLife, 4:e07192, July 2015.

[17] Jonathan Paul Newman, Riley Zeller-Townson, Ming-fai Fong, Sharanya Arcot Desai, Robert E. Gross, and Steve M. Potter. Closed-Loop, Multichannel Experimentation Using the Open-Source NeuroRighter Electrophysiology Platform. Frontiers in Neural Circuits, 6, 2013.

[18] OpenPCR - the $499 Open Source PCR Machine / Thermal Cycler.

[19] Bas Wijnen, Emily J. Hunt, Gerald C. Anzalone, and Joshua M. Pearce. Open-Source Syringe Pump Library. PLOS ONE, 9(9):e107216, September 2014.

[20] Chris N. Takahashi, Aaron W. Miller, Felix Ekness, Maitreya J. Dunham, and Eric Klavins. A low cost, customizable turbidostat for use in synthetic circuit characterization. ACS synthetic biology, 4(1):32–38, January 2015.

[21] Stefan A. Hoffmann, Christian Wohltat, Kristian M. Müller, and Katja M. Arndt. A user-friendly, low-cost turbidostat with versatile growth rate estimation based on an extended Kalman lter. PLOS ONE, 12(7):e0181923, July 2017.

[22] Brandon G. Wong, Christopher P. Mancuso, Szilvia Kiriakov, Caleb J. Bashor, and Ahmad S. Khalil. Precise, automated control of conditions for high-throughput growth of yeast and bacteria with eVOLVER. Nature Biotechnology, 36(7):614–623, 2018.

[23] Andre Maia Chagas, Lucia L. Prieto-Godino, Aristides B. Arrenberg, and Tom Baden. The 100-euro lab: A 3d-printable open-source platform for fluorescence microscopy, optogenetics, and accurate temperature control during behaviour of zebrafish, Drosophila, and Caenorhabditis elegans. PLOS Biology, 15(7):e2002702, July 2017.

[24] G. Gürkan and K. Gürkan. Incu-Stream 1.0: An Open-Hardware Live-Cell Imaging System Based on Inverted Bright-Field Microscopy and Automated Mechanical Scanning for Real-Time and Long-Term Imaging of Microplates in Incubator. IEEE Access, 7:58764–58779, 2019.

[25] Hongquan Li, Hazel Soto-Montoya, Maxime Voisin, Lucas Fuentes Valenzuela, and Manu Prakash. Octopi: Open configurable high-throughput imaging platform for infectious disease diagnosis in the field. bioRxiv, page 684423, June 2019.

[26] Tomas Aidukas, Regina Eckert, Andrew R. Harvey, Laura Waller, and Pavan C. Konda. Low-cost, sub-micron resolution, wide-field computational microscopy using opensource hardware. Scientific Reports, 9(1):7457, May 2019.

[27] James P. Sharkey, Darryl C. W. Foo, Alexandre Kabla, Jeremy J. Baumberg, and Richard W. Bowman. A one-piece 3d printed flexure translation stage for open-source microscopy. Review of Scientific Instruments, 87(2):025104, February 2016.

[28] Karol P. Szymula, Michael S. Magaraci, Michael Patterson, Andrew Clark, Sevile G. Mannickarottu, and Brian Y. Chow. An Open-Source Plate Reader. Biochemistry, 58(6):468–473, February 2019.

[29] Vasco Ribeiro Pereira and Bill Stephen Hosker. Low-cost (5-euro), open-source, potential alternative to commercial spectrophotometers. PLOS Biology, 17(6):e3000321, June 2019.

[30] Gaurav Byagathvalli, Soham Sinha, Yan Zhang, Mark P. Styczynski, Janet Standeven, and M. Saad Bhamla. ElectroPen: An ultralow-cost piezoelectric electroporator. bioRxiv, page 448977, December 2018.

[31] Andrew P. Hill, Peter Prince, Evelyn Pia Covarrubias, C. Patrick Doncaster, Jake L. Snaddon, and Alex Rogers. AudioMoth: Evaluation of a smart open acoustic device for monitoring biodiversity and the environment. Methods in Ecology and Evolution, 9(5):1199–1211, 2018.

[32] Texas Instruments. AN-1515 A comprehensive Study of the Howland Current Pump. Application report SNOA474A, 2008.

[33] Wayne Stambaugh, Jean-Pierre Charras, and Dick Hollenbeck. KiCad EDA.

[34] Joshua I. Sanders and Adam Kepecs. A low-cost programmable pulse generator for physiology and behavior. Frontiers in Neuroengineering, 7, 2014.

[35] James Olds and Peter Milner. Positive reinforcement produced by electrical stimulation of septal area and other regions of rat brain. Journal of Comparative and Physiological Psychology, 47(6):419–427, 1954.

[36] Ilana B. Witten, Elizabeth E. Steinberg, Soo Yeun Lee, Thomas J. Davidson, Kelly A. Zalocusky, Matthew Brodsky, Ofer Yizhar, Saemi L. Cho, Shiaoching Gong, Charu Ramakrishnan, Garret D. Stuber, Kay M. Tye, Patricia H. Janak, and Karl Deisseroth. Recombinase-Driver Rat Lines: Tools, Techniques, and Optogenetic Application to Dopamine-Mediated Reinforcement. Neuron, 72(5):721–733, December 2011.

[37] Elizabeth E. Steinberg, Josiah R. Boivin, Benjamin T. Saunders, Ilana B. Witten, Karl Deisseroth, and Patricia H. Janak. Positive reinforcement mediated by midbrain dopamine neurons requires d1 and d2 receptor activation in the nucleus accumbens. PLOS ONE, 9(4):1–9, 04 2014.

[38] Xiao Han, Yi He, Guo-Hua Bi, Hai-Ying Zhang, Rui Song, Qing-Rong Liu, Josephine M. Egan, Eliot L. Gardner, Jing Li, and Zheng-Xiong Xi. Cb1 receptor activation on vglut2-expressing glutamatergic neurons underlies delta-9-tetrahydrocannabinol (delta-9-thc)-induced aversive effects in mice. Scientific Reports, 7(1), September 2017.

[39] William A. Carlezon and Elena H. Chartoff. Intracranial self-stimulation (ICSS) in rodents to study the neurobiology of motivation. Nature Protocols, 2(11):2987–2995, 2007. 8

