## Supplementary material for "Stimjim: open source hardware for precise electrical stimulation": Stimjim circuit schematic

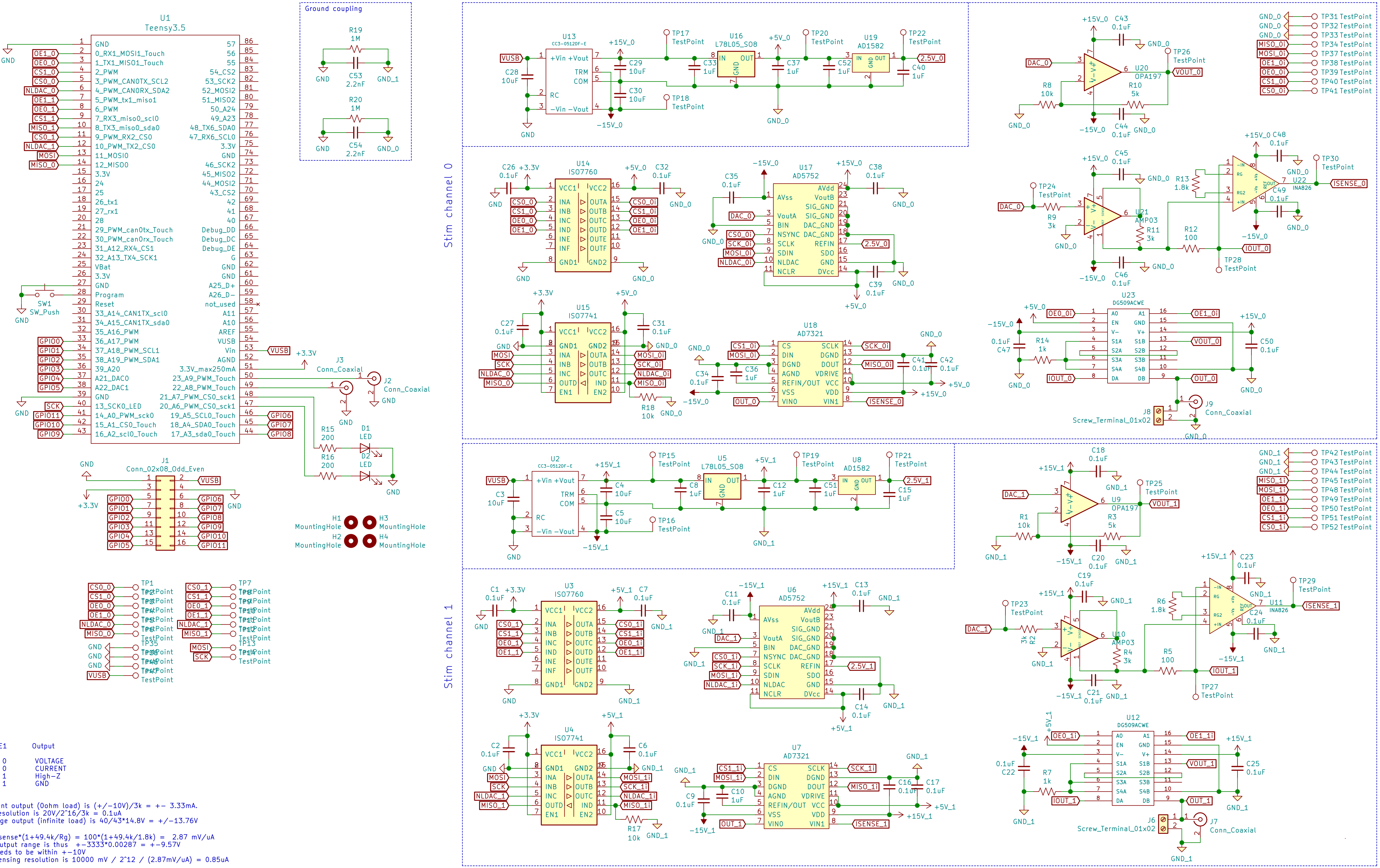


| OE0 | OE1 | Output |
|---|---|---|
| 0 | 0 | VOLTAGE |
| 1 | 0 | CURRENT |
| 0 | 1 | High-Z |
| 1 | 1 | GND |

Max current output (0ohm load) is (+/-10V)/3k = +- 3.33mA.
Current resolution is 20V/2^16/3k = 0.1uA
Max voltage output (infinite load) is 40/43*14.8V = +/-13.76V

gain is Rsense*(1+49.4k/Rg) = 100*(1+49.4k/1.8k) = 2.87 mV/uA
Voltage output range is thus +-3333*0.00287 = +-9.57V
output needs to be within +-10V
current sensing resolution is 10000 mV / 2^12 / (2.87mV/uA) = 0.85uA
